# MDM4 is an essential disease driver targeted by 1q gain in Burkitt lymphoma

**DOI:** 10.1101/289363

**Authors:** Jennifer Hüllein, Mikołaj Słabicki, Maciej Rosolowski, Alexander Jethwa, Stefan Habringer, Katarzyna Tomska, Roma Kurilov, Junyan Lu, Sebastian Scheinost, Rabea Wagener, Zhiqin Huang, Marina Lukas, Olena Yavorska, Hanne Helferich, René Scholtysik, Kyle Bonneau, Donato Tedesco, Ralf Küppers, Wolfram Klapper, Christiane Pott, Stephan Stilgenbauer, Birgit Burkhardt, Markus Löffler, Lorenz Trümper, Michael Hummel, Benedikt Brors, Marc Zapatka, Reiner Siebert, consortium MMML, Ulrich Keller, Wolfgang Huber, Markus Kreuz, Thorsten Zenz

## Abstract

Oncogenic MYC activation promotes cellular proliferation in Burkitt lymphoma (BL), but also induces cell cycle arrest and apoptosis mediated by *TP53*, a tumor suppressor gene that is mutated in 40% of BL cases. To identify therapeutic targets in BL, we investigated molecular dependencies in BL cell lines using RNAi-based, loss-of-function screening. By integrating genotypic and RNAi data, we identified a number of genotype-specific dependencies including the dependence of *TCF3/ID3* mutant cell lines on TCF3 and of *MYD88* mutant cell lines on TLR signaling. *TP53* wild-type (TP53wt) BL were dependent on MDM4, a negative regulator of TP53. In BL cell lines, MDM4 knockdown induced cell cycle arrest and decreased tumor growth in a xenograft model in a p53-dependent manner, while small molecule inhibition of the MDM4-p53 interaction restored p53 activity resulting in cell cycle arrest. Consistent with the pathogenic effect of MDM4 upregulation in BL, we found that TP53wt BL samples were enriched for gain of chromosome 1q which includes the *MDM4* locus. 1q gain was also enriched across non-BL cancer cell lines (n=789) without *TP53* mutation (23% in TP53wt and 12% in TP53mut, p<0.001). In a set of 216 cell lines representing 19 cancer entities from the Achilles project, MDM4 was the strongest genetic dependency in TP53wt cell lines (p<0.001).

Our findings show that in TP53wt BL, MDM4-mediated inhibition of p53 is a mechanism to evade cell cycle arrest. The data highlight the critical role of p53 as a tumor suppressor in BL, and identifies MDM4 as a key functional target of 1q gain in a wide range of cancers, which is therapeutically targetable.

## Introduction

Burkitt lymphoma (BL) is an aggressive B cell lymphoma that is characterized by translocation of the *MYC* gene to immunoglobulin loci (Molyneux et al., 2012). While oncogenic MYC promotes cell growth and proliferation (Dalla-Favera et al., 1982; Taub et al., 1982), it also evokes failsafe mechanisms such as p53 activation that have to be overcome for transformation (Evan et al., 1992; Meyer et al., 2006). About 30% of BL acquire *TP53* mutations and as a consequence cells do not respond to apoptotic stimuli (Bhatia et al., 1992; Gaidano et al., 1991; O’Connor et al., 1993). As shown in mice, mutations in the conserved *Myc* box I prevent the induction of apoptosis via Bim as an alternative mechanism to *TP53* mutation, providing further evidence that secondary lesions cooperate with oncogenic MYC induction (Hemann et al., 2005).

Recent mutational cartography efforts in BL identified additional recurrent mutations in *TCF3*, *ID3*, *GNA13*, *RET*, *PIK3R1*, *DDX3X*, *FBXO11*, and the SWI/SNF genes *ARID1A* and *SMARCA4* (Kretzmer et al., 2015; Love et al., 2012; Richter et al., 2012; Schmitz et al., 2012). BL also display copy number alterations (CNAs) in addition to the *MYC* translocation (Scholtysik et al., 2010), targeting chromosomes 1q, 3p27, 13q31, 17p13 (including *TP53*) and 9p21.2 (including *CDKN2A*). A gain of 1q is found in 30% of BL and often affects large regions (Salaverria et al., 2008), which has contributed to the limited understanding of oncogenic mechanisms involved. The clinical implications of these mutations and CNAs are currently unclear.

RNAi-based genomics screens allow querying of functional dependencies in an unbiased fashion and in high-throughput (Mohr et al., 2014). Using panels of representative cell lines, context-specific vulnerabilities have been linked to genetic and pathological subgroups (Cheung et al., 2011). The Achilles Project reported comprehensive screening data in 501 cell lines using RNAi (Cowley et al., 2014; Tsherniak et al., 2017). While activating mutations caused direct oncogene addiction, as seen in cell lines with *BRAF*, *KRAS* or *PI3K* mutation, secondary gene dependencies were observed for loss-of-function mutations in tumor suppressor genes, such as *ARID1A* (Helming et al., 2014). Integration of gene expression and drug sensitivity profiles may provide further insight into the molecular basis of diseases and might be used to tailor targeted therapies (Marcotte et al., 2016).

For a comprehensive dissection of molecular dependencies in BL, we performed a loss-of-function RNAi screen across a panel of genetically characterized BL cell lines and intersected our findings on genotype-specific essential genes with the genetic profile of a well-annotated patient cohort.

## Results

### Landscape of essential genes in BL

To identify therapeutic targets in BL, we investigated molecular dependencies in BL cell lines using RNAi-based loss-of-function screening. We used 27,495 shRNAs to silence 5,045 genes and assessed changes in shRNA abundance after culturing the cells for two weeks (Figure 1A). On average 24% of shRNAs were depleted at least two-fold and shRNAs targeting core essential complexes, including the ribosome and the proteasome, were specifically lost (68% and 47%, respectively) (Figure 1B). To evaluate the viability effect of individual gene knock-downs, we calculated weighted z-scores that combine the effect of shRNAs targeting the same gene and emphasize strong fold-changes (Dai et al., 2014; Kim and Tan, 2012) (see Methods). Common essential genes, as defined on the basis of previous RNAi screens (Hart et al., 2014), showed significantly lower scores compared to non-essential genes (p<0.001, Figure 1C). Notably, while a subset of genes was essential in all cell lines, we also observed cell line specific viability effects (Figure S1A, B).

**Figure 1.**
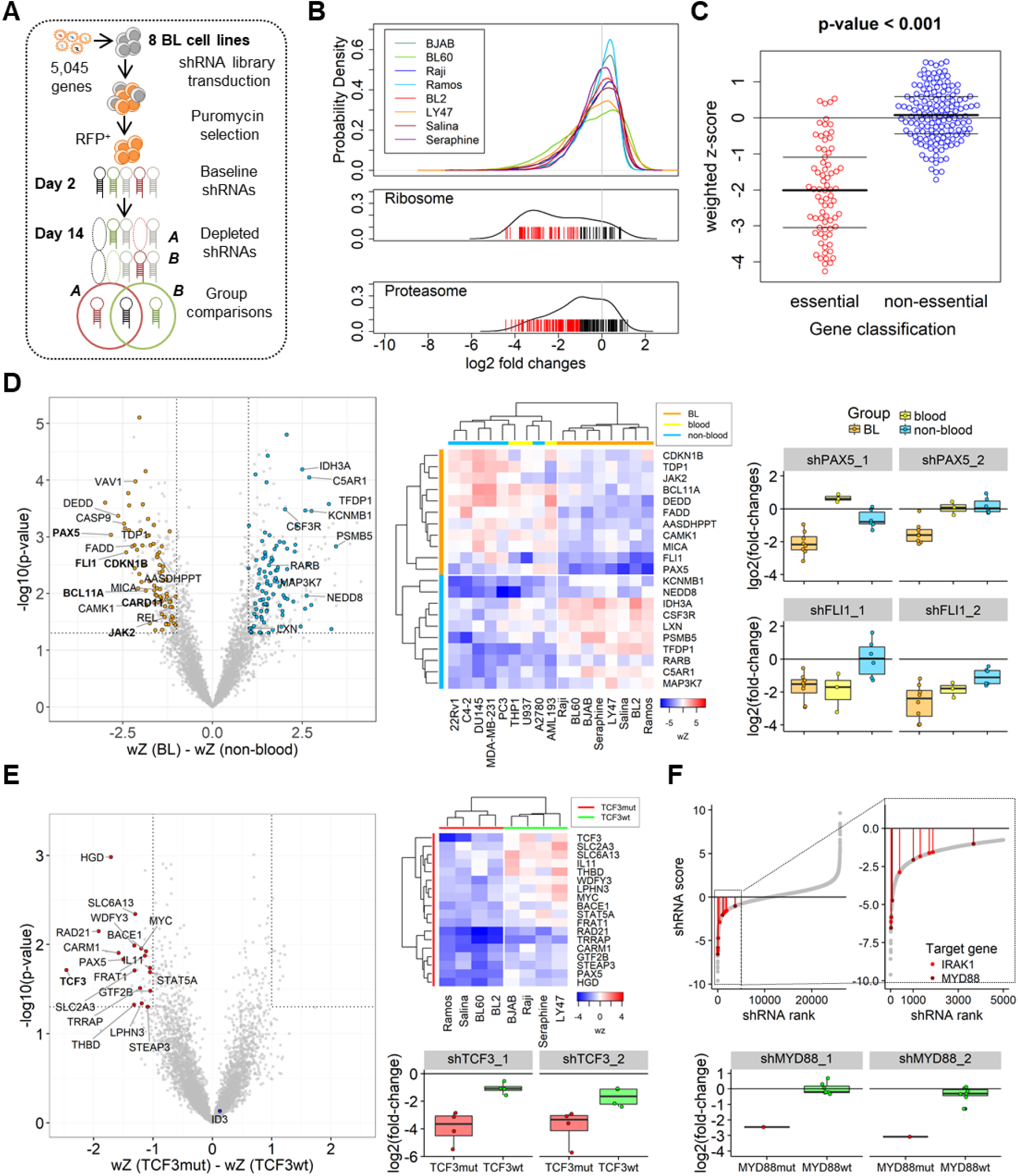
RNAi screening reveals context-specific vulnerabilities in BL. **(A)** Layout of the RNAi screen for the identification of viability genes in BL cell lines. After lentiviral transfer of a pooled shRNA library, shRNA abundance was determined by high-throughput sequencing. shRNAs interfering with cell growth or proliferation were lost over time. **(B)** shRNA depletion after two weeks of culture. The upper plot shows changes in shRNA abundance per cell line. The lower plots show mean fold-changes of shRNAs targeting the ribosome or proteasome, respectively. shRNAs with a fold-change of 2 or lower are marked in red, indicating specific depletion of shRNAs targeting core cellular complexes. **(C)** The toxicity of a gene knock-down was measured in weighted z-scores (wZ) that combine the effect of single shRNAs targeting the same gene. The mean wZ-score is shown for common essential genes (n=73) and non-essential genes (n=149) as defined previously (Hart et al., 2004). **(D)** Essential genes in BL (orange) were compared to essential genes in six solid cancer cell lines (MDA-MB-231, A2780, C4-2, R22v1, PC3, DU-145) (blue). The volcano plot shows differences in wZ-scores and the rectangles mark the cut-off values at a p-value of 0.05 and difference of mean wZ-score of 1. The ten strongest classifiers of BL or non-blood cancer cell lines, respectively, based on differential shRNA fold-changes are labeled. The heatmap shows the wZ-scores of the ten highest ranking genes and the B-cell marker PAX5 (rank 37). wZ-scores in two AML and one DLBCL cell lines (yellow) are shown to differentiate between BL- and lineage-specific essential genes. shRNA fold-changes are shown for a BL-specific essential gene (*PAX5*) and a blood-lineage specific gene (*FLI1*). **(E)** Essential genes in four cell lines with *TCF3* and/or *ID3* mutation were compared to four wild-type cell lines. Genes with genotype-specific viability effects were selected as described above and are highlighted in the volcano plot and displayed in the heatmap of wZ-scores. The boxplots show the specific loss of shRNAs targeting *TCF3* in the presence of *TCF3*-activating mutations. **(F)** shRNAs were ranked by their differential effects in BL-2 (MYD88mut) and seven MYD88wt BL cell lines. shRNAs targeting *MYD88* or its downstream target *IRAK1* were specifically lost in the mutant context.

To investigate essential genes in the context of BL, we probed our data against RNAi screening results using the same set of shRNAs in six carcinoma cell lines (C4-2, DU145, PC3, R22v1, MDA-MB-231, A2780) and three cell lines of myeloid and lymphoid origin (AML193, THP1, U937) (Figure S1C, Table S1). We ranked shRNAs based on their differential effects between two cell line groups and calculated a gene classification score as a measurement of their strength to distinguish between the groups (Cheung et al., 2011) (Table S2). We then selected genes that were predictors of an entity group and showed strong differential viability effects based on the weighted z-scores. Using this method, we identified 76 genes essential in BL, including genes associated with hematopoietic cell differentiation (*FLI1*, *BCL11A*) or B cell development and activation (*PAX5*, *CDKN1B*, *JAK2*, *CARD11*) (Figure 1D, *left*). The transcription factor FLI1 is a key regulator of the hematopoietic system and B cell development (Zhang et al., 2008; Zochodne et al., 2000). In our screen, knock-down of FLI1 was also toxic to blood-lineage derived non-BL cell lines, while PAX5, a marker of early B-cell development, was an essential gene exclusively in BL (Figure 1D, *middle*/*right*).

### Genotype-specific dependencies in BL

To identify exploitable vulnerabilities in the context of gene mutations, we performed RNA sequencing of the BL cell lines included in the RNAi screen. Genes that are recurrently mutated in BL, including *TP53*, *ID3*, *TCF3*, *DDX3X*, *FOXO1* and *GNA13* (Kretzmer et al., 2015; Love et al., 2012; Richter et al., 2012; Schmitz et al., 2012), were represented by our cell line models (Table S3). We therefore investigated essential genes in the respective genotype groups. Mutations in the transcription factor *TCF3* lead to oncogene activation and loss-of-function mutations of its inhibitor *ID3* are often observed as a complementary mechanism of TCF3 activation (Schmitz et al., 2012). Therefore, cell lines carrying either *TCF3* or *ID3* mutation were treated as one group. The four cell lines with *TCF3*/*ID3* mutation were strongly dependent on TCF3 expression, indicating oncogene addiction (p<0.01) (Figure 1E). In line with the loss of function effect of mutations in *ID3*, ID3 silencing was not toxic (Figure 1E, *left*). The cell line BL-2 harbors the activating p.S219C mutation in MYD88, an adaptor protein involved in Toll-Like-Receptor signaling and NF-kB activation. Among the shRNAs that were toxic in the MYD88mut context, we identified an enrichment for shRNAs targeting MYD88 or its direct downstream mediator IRAK1 (Figure 1F). Encouraged by the ability to uncover oncogene addiction, we expanded our analysis of genotype-specific vulnerabilities to *DDX3X*, *FOXO1*, *GNA13* and *TP53* (Table S2; Figure S1D). *TP53* mutation was associated with the strongest differential viability effects (gene classification scores > 2) and we therefore focused on *TP53*-specific vulnerabilities (Figure 2A, *left*).

**Figure 2.**
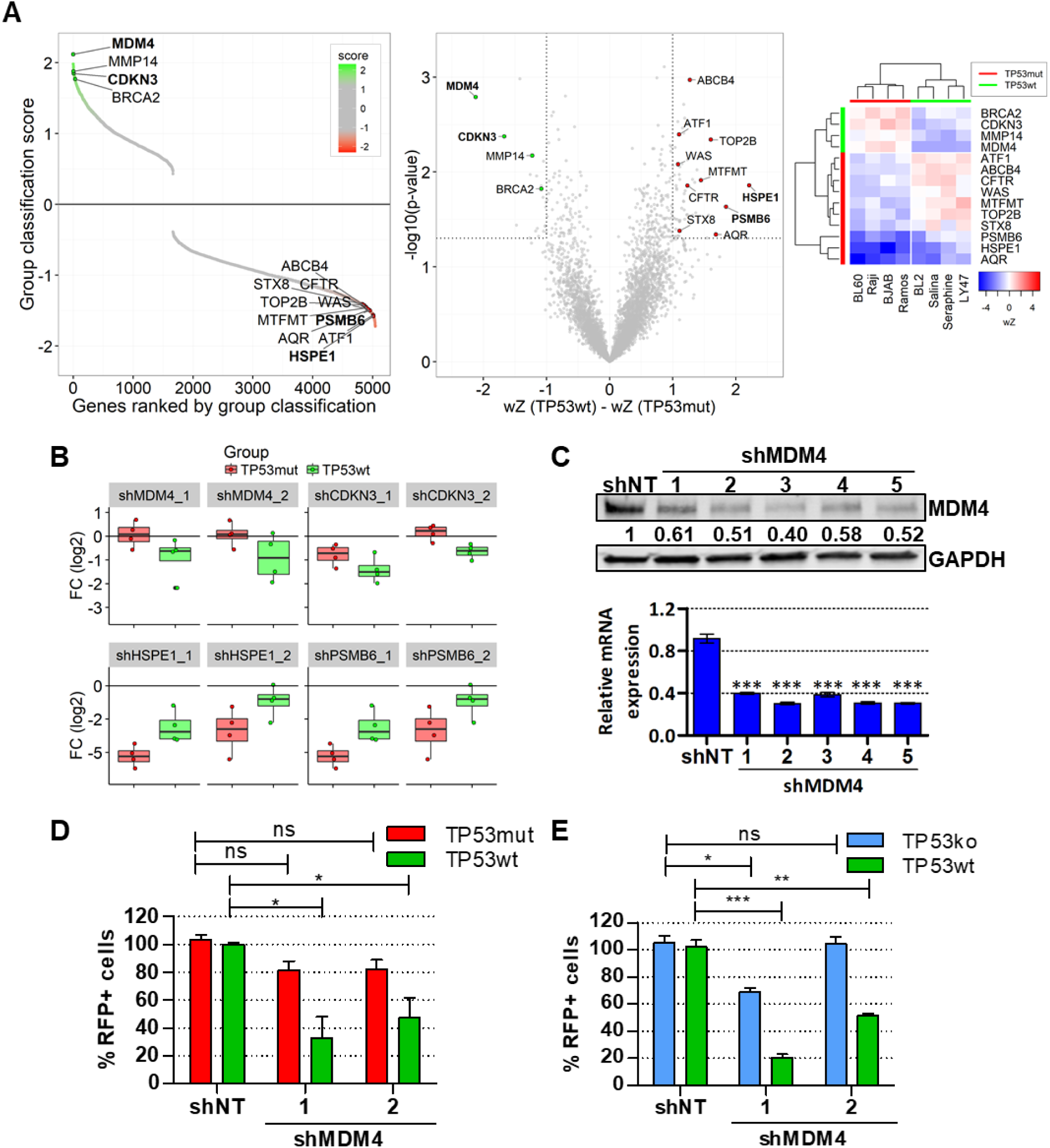
MDM4 is essential in TP53wt BL. **(A)** (*left*) Genes were ranked from genes with lower shRNA counts in the TP53wt group to lower counts in the TP53mut group. The gene score expresses the strength of the gene in distinguishing between the groups. (*middle*) The volcano plot shows differences in wZ-scores between four TP53wt and four TP53mut BL cell lines as described for Figure 1D. (*right*) wZ-scores of genes with *TP53*-status dependent viability effects. **(B)** shRNA fold-changes from the RNAi screen for candidate genes essential in TP53wt (*MDM4*, *CDKN3*) or TP53mut (*HSPE1*, *PSMB6*) BL cell lines. **(C)** RT-qPCR and immunoblot for MDM4 level five days after shRNA delivery. Expression values were normalized to GAPDH and non-targeting shRNA. RT-qPCR error bars indicate the mean with standard deviation of triplicate measurements (***: p<0.001), for immunoblot, normalized ratio of protein intensity is indicated. **(D, E)** Growth competition assay for two independent shRNAs targeting MDM4. Four TP53wt, eight TP53mut and one TP53ko cell line were transduced with shNT, shMDM4_1 or shMDM4_2 at a rate of 50% and the fraction of shRNA expressing cells was monitored over 14 days by co-expression of RFP. The proportion of RFP+ cells 14 days after transduction was normalized to day 3. **(D)** Error bars in the comparison of TP53wt and TP53mut cell lines indicate results across cell lines using the mean of two independent experiments +/-SEM. **(E)** Error bars in the comparison of the isogenic Seraphine TP53wt and TP53ko cell line indicate triplicate measurements (*: p<0.05; **: p<0.01; ***: p<0.001).

### p53 pathway in BL

Based on shRNA depletion in four *TP53* mutant (TP53mut) versus four *TP53* wild-type (TP53wt) cell lines, we identified viability genes specific to both groups (Figure 2A). TP53mut cell lines were more sensitive to depletion of *PSMB6* (Figure 2A, B) and additional proteasomal subunits (Figure S2). A gain-of-function phenotype of mutant p53 has been associated with increased proteasome activity (Walerych et al., 2016) and proteasome inhibition was shown to reduce mutant p53 expression (Halasi et al., 2014). We also identified the heat shock protein *HSPE1* as an essential gene in mutant cell lines (Figure 2A, B). Upon cellular stress, heat shock proteins regulate the stability of wild-type and mutant p53 and HSP90 inhibitors were shown to be effective in p53 mutant cell lines (Blagosklonny et al., 1996; Li et al., 2011; Muller et al., 2008).

Genes essential in TP53wt cell lines included the cell cycle regulators *CDKN3* (Figure 2A, B), a spindle checkpoint phosphatase activated by p21 (Baldi et al., 2011). The strongest gene dependency of the TP53wt group was *MDM4* (gene score=2.12; Figure 2A, *left*) that inactivates p53-mediated transcription by blocking of its transactivation domain (Shvarts et al., 1997). Notably, as Epstein-Barr virus (EBV) associated proteins deregulate cell cycle checkpoints and quench the p53 pathway by deubiquitination of the p53 inhibitor MDM2 (Fish et al., 2017; Saha et al., 2009), we confirmed a balanced distribution of EBV infection status among TP53wt and TP53mut BL cell lines (Table S3).

### MDM4 is essential in TP53wt BL

shRNAs targeting MDM4 were more toxic in TP53wt than TP53mut cell lines (Figure 2B) and all shRNAs efficiently reduced MDM4 mRNA and protein levels (Figure 2C). Using two non-overlapping shRNAs, we validated the screen findings in a growth competition assay in all BL cell lines included in the RNAi screen and four additional TP53mut cell lines. shRNAs were co-expressed with red fluorescent protein (RFP) in ~50% of cells and the fraction of RFP+/shRNA+ cells was monitored over time. TP53wt cell lines (n=4) showed a significant reduction of cells with MDM4 knock-down, which was not observed for TP53mut cell lines (n=8), confirming the screen results (Figure 2D).

To further test whether the observed effects were dependent on p53, we generated a p53 knock-out cell line based on the TP53wt cell line Seraphine (Figure S3). Seraphine-TP53ko showed a decreased basal apoptotic rate, while the cell cycle progression was unaffected (Figure 3B, S3B). When we silenced MDM4 in Seraphine-TP53wt, we observed significantly stronger loss of RFP+ cells compared to Seraphine-TP53ko, confirming that the viability effects following MDM4 depletion were p53-dependent (Figure 2E).

**Figure 3.**
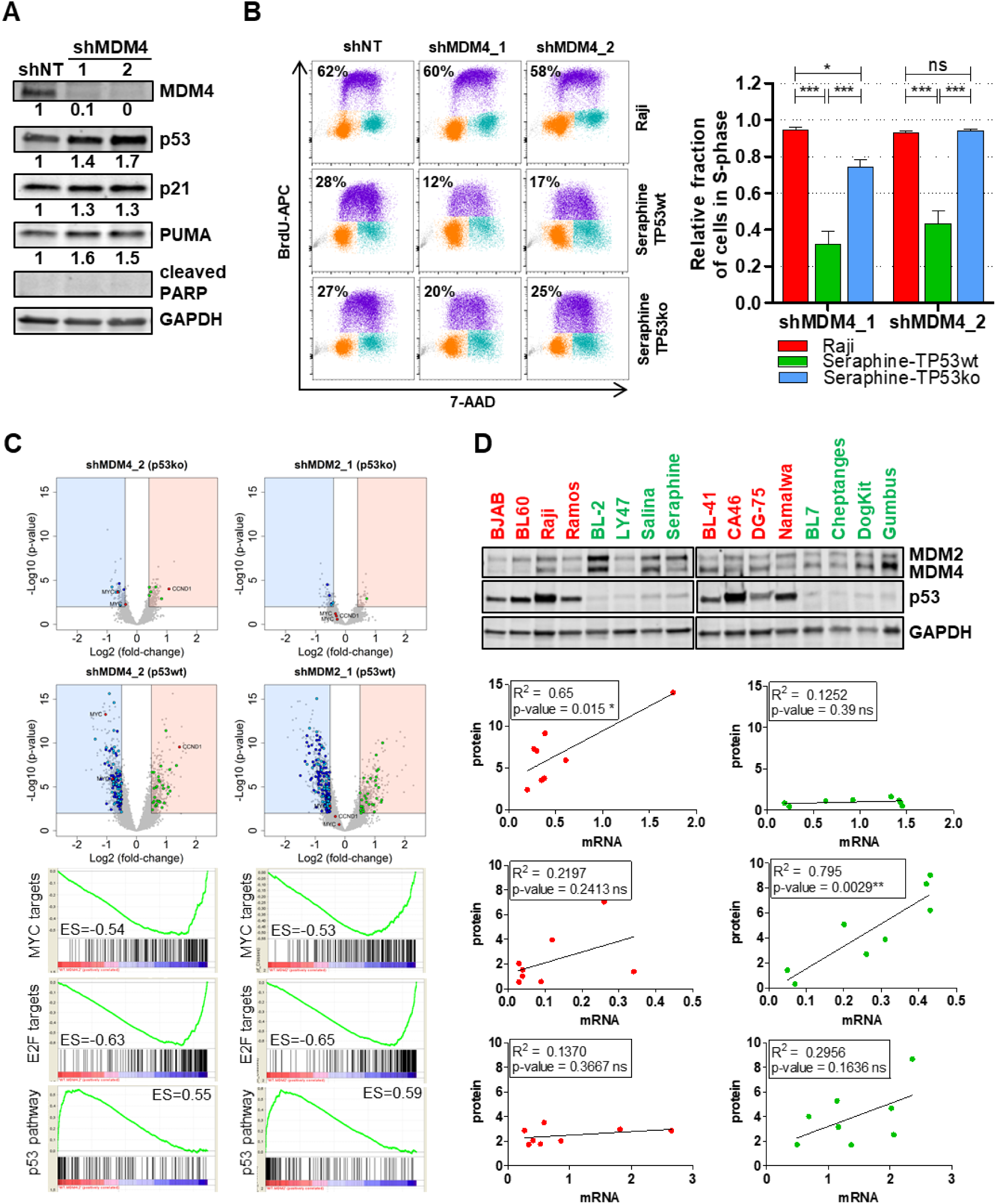
MDM4 depletion reactivates p53 and induces G1 arrest. **(A)** Immunoblot analysis of p53, p21, PUMA, and cleaved PARP was performed in a representative TP53wt cell line (Seraphine) after MDM4 knock-down with two independent shRNAs. Protein lysates were analyzed on day 5 post-transduction. Band intensities were normalized to GAPDH and shNT. Silencing of MDM4 led to p53 and p21 induction without PARP cleavage. **(B)** Analysis of the cell cycle profile after MDM4 knock-down by flow cytometry. Cycling cells were determined by BrdU incorporation and total DNA was stained with 7-AAD. The plots show a representative measurement. Quantification of triplicate experiments is shown on the right (ns: p>0.05, *: p <0.05, ***: p <0.001). **(C)** Global gene expression changes after MDM4 and MDM2 knock-down in isogenic Seraphine cell lines. Expression levels were normalized to shNT and gene set enrichment analysis was performed in the TP53wt cell line using the java GSEA software (http://software.broadinstitute.org/gsea/downloads.jsp). Enrichment curves for the most enriched pathways are shown below the volcano plots. Genes from suppressed pathways are highlighted in blue and genes from enriched pathways in green. Genes highlighted in red were changed after MDM4, but not after MDM2 knock-down (cut-off –log10(p-value) > 2, log2(fold-change) < -0.5 or > 0.5). **(D)** Immunoblot and RT-qPCR analysis of basal expression levels for MDM4, MDM2 and p53 in eight TP53wt (green) and eight TP53mut (red) BL cell lines. Shown here is the Pearson correlation coefficient (R²).

### MDM4 promotes cell cycle progression by p53 inactivation

To understand the downstream effects of MDM4 depletion in BL, we assessed protein levels of p53 and known p53 targets. MDM4 knock-down in TP53wt cells increased p53 protein level and induced the pro-apoptotic Bcl-2 family member PUMA and the cell cycle inhibitor p21 (Figure 3A). Since MDM4 downregulation did not cause apoptosis as determined by absence of PARP cleavage (Figure 3A), we analyzed the cell cycle profile in the presence or absence of functional p53 after MDM4 silencing. In the TP53wt context, shRNAs targeting MDM4 decreased cycling cells compared to a non-targeting shRNA (shNT, p<0.001), which was not observed in the TP53mut or the TP53ko cell line (Figure 3B).

We next determined global gene expression changes after MDM4 and MDM2 silencing in the TP53wt and TP53ko Seraphine cell lines (Figure 3C, Table S4). Silencing of MDM4 or MDM2 induced strong changes only in the presence of p53 and affected similar pathways. Using gene set enrichment analysis for cancer hallmark genes (MSigDB), we identified p53 targets as the strongest upregulated pathway, while prominent survival and proliferation pathways, including MYC and E2F targets, were downregulated. This suggests that most effects were mediated by p53 activation, in accordance with a previous report on genes commonly regulated after MDM4 or MDM2 knock-down (Heminger et al., 2009). We also compared genes differentially regulated by MDM2 or MDM4 silencing (Figure S4). Downregulation of *MYC* and upregulation of *CCND1* were exclusively seen after MDM4 knock-down, indicating potential differences in pathway contribution exerted by MDM4 over MDM2 (Figure 3C, S4). We next examined the basal protein and mRNA expression levels of p53, MDM4 and MDM2 in a panel of BL models (Figure 3D). p53 protein was detected at higher level in all TP53mut cell lines (p<0.01) as described previously (Bartek et al., 1991; Farrell et al., 1991), while p53 mRNA levels were lower. While wild-type p53 is rapidly turned-over in a negative feed-back loop mediated by MDM2, mutant p53 protein accumulates as a result of disrupted proteasomal decay pathways (Vijayakumaran et al., 2015). MDM4 mRNA was significantly higher in TP53wt BL cell lines (p=0.05) and was correlated with protein expression (p<0.01) (Figure 3D). MDM2 levels were also elevated in TP53wt cell lines, but not significantly (Figure 3D).

### MDM4 is a therapeutic target in TP53wt BL

To evaluate the potential of MDM4 as a therapeutic target in TP53wt BL *in vivo*, we determined the effect of MDM4 silencing on tumor growth in a mouse xenograft model. After transduction, cell lines representing TP53wt (Seraphine), TP53ko (Serphine-TP53ko) and TP53mut (Raji) were injected subcutaneously into the flanks of immunodeficient mice (Herhaus et al., 2016). To quantify tumor formation and dynamic growth, we measured fludeoxyglucose (FDG) uptake in positron emission tomography (PET). *In vivo* tumor formation was significantly reduced after MDM4 knockdown in the presence of wild-type p53 (p<0.05) (Figure 4A, B).

**Figure 4.**
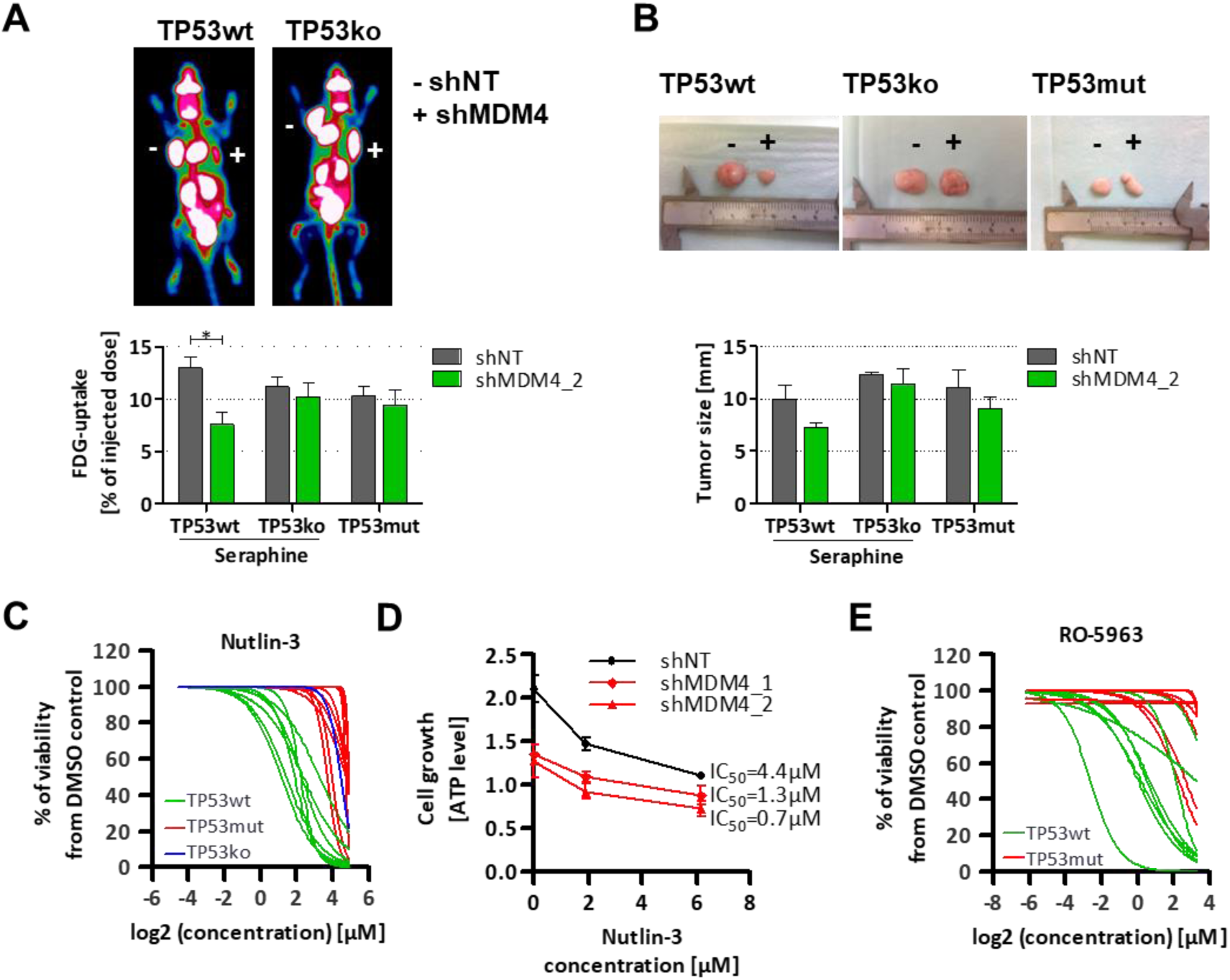
MDM4 is a therapeutic target in TP53wt BL. **(A, B)** MDM4 depletion reduces tumor growth in a mouse xenograft model. Indicated cell lines expressing shNT or shMDM4 were subcutaneously injected into the left (shNT) or right (shMDM4) flank of immunodeficient mice. **(A)** Exemplary images from FDG-PET analysis and quantification of FDG-uptake and **(B)** excised xenografts are shown. Error bars indicate mean of three mice per cell line and shRNA construct with standard error. **(C)** Cell line sensitivity to Nutlin-3 stratified by *TP53* mutation. Ten TP53mut, seven TP53wt and one TP53ko BL cell line were exposed to Nutlin-3 for 48h and viability was assayed by ATP content and normalized to DMSO treated samples. **(D)** Cell proliferation after combined inhibition of MDM4 and MDM2 in Seraphine-TP53wt. Cells expressing shNT, shMDM4_1 or shMDM4_2 were incubated for two days in the presence of Nutlin-3. Cell content was determined by ATP content. Error bars indicate the average over two independent experiments with standard deviation. **(E)** Cell line viability of seven TP53mut and seven TP53wt BL cell lines exposed to the dual MDM2/MDM4 inhibitor RO-5963 for 48h. Viability was assayed by ATP content and normalized to DMSO treated samples.

Restoration of p53 activity is an attractive therapeutic approach for treatment of cancer (Burgess et al., 2016). The small molecule inhibitor Nutlin-3 targets the p53 inhibitor MDM2 and therefore restores signaling through the p53 pathway (Vassilev et al., 2004). Nutlin-3 increased intracellular p53 level and induced apoptosis in Seraphine-TP53wt, but not in Seraphine-TP53-ko (Figure S3A). All TP53wt cell lines were sensitive to Nutlin-3 (Figure 4C). Despite the high sequence homology of MDM2 and MDM4, Nutlin-3 targets MDM2 with a much higher binding affinity (Patton et al., 2006). Moreover, overexpression of MDM4 can lead to resistance against MDM2-targeting drugs (Hu et al., 2006; Patton et al., 2006). We therefore tested the potential of combined inhibition of MDM4 and MDM2 in BL. MDM4 knock-down increased cytotoxic effects of Nutlin-3 (Figure 4D). TP53wt cell lines were sensitive towards the dual-specificity inhibitor RO-5963, that targets MDM2 and MDM4 (Graves et al., 2012) (Figure 4E). This data provides a rational for targeting MDM4/2 in TP53wt BL.

### Gain of MDM4 on chr1q provides an alternative to *TP53* mutations in BL

To understand the role of the p53 pathway in BL, we analyzed the genetic profile of aggressive B-cell lymphomas stratified into BL, diffuse large B cell lymphoma (DLBCL) or cases with intermediate phenotype (Table S5). *TP53* mutations were found in 28/61 (45.9%) of BL samples and were significantly more frequent in BL than in DLBCL (p<0.001) (Figure 5A). *MYC* box I mutations were previously reported to be mutually exclusive with *TP53* mutations and to serve as an alternative mechanism to escape apoptotic pathways in the presence of wild-type *TP53* (Hemann et al., 2005). *MYC* mutations were present in 37/56 BL samples (66.1%) and the *MYC* box I residues 56-58 were affected in 20 (35.7%) cases (Figure 5B). Notably, *MYC* box I mutations frequently co-occurred with *TP53* mutations (Figure 5B).

**Figure 5.**
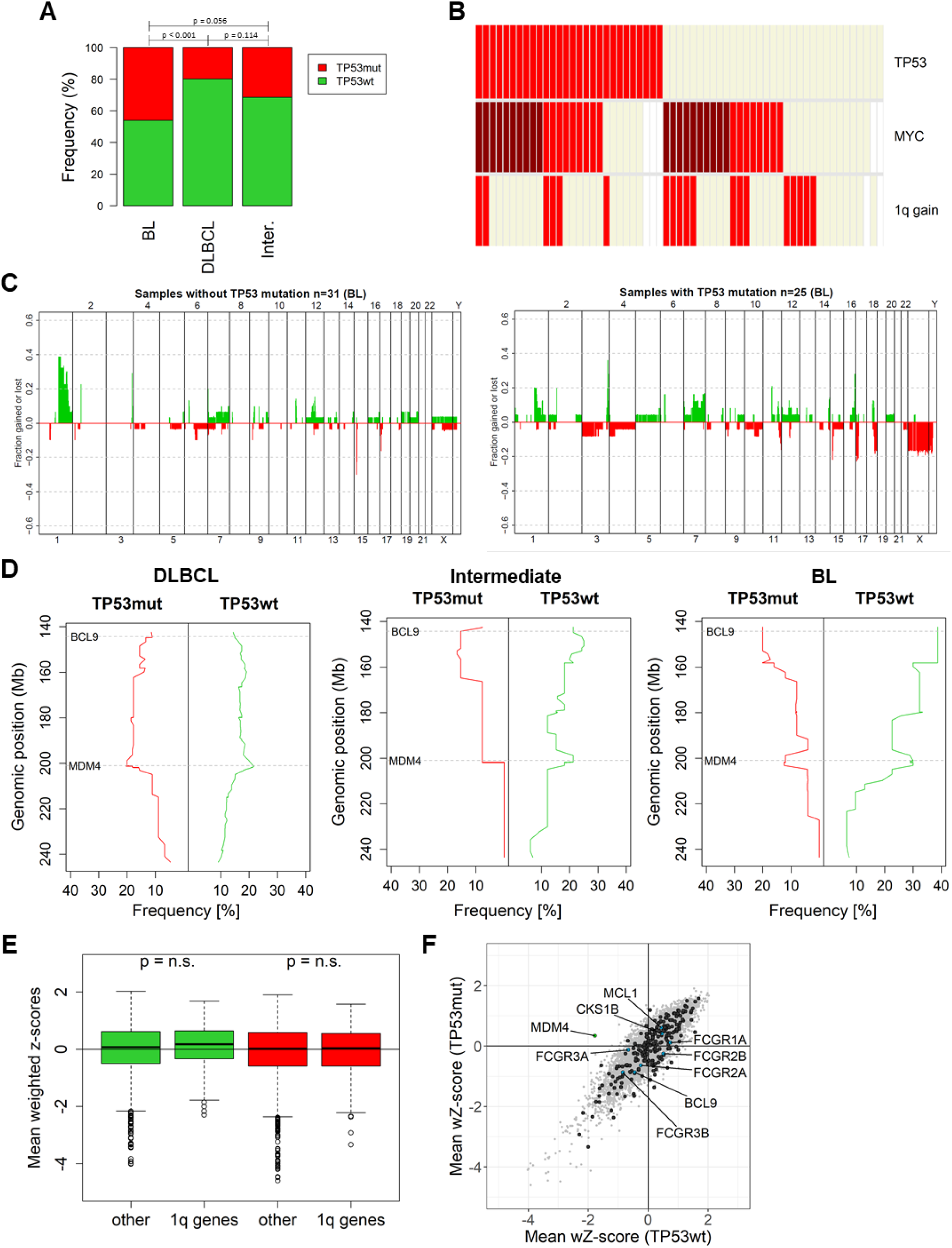
Genetic aberrations frequently affect the p53 pathway in BL. **(A)** Incidence of *TP53* mutations in BL (n=61), DLBCL (n=297) and the “intermediate” group based on gene expression (n=54) as determined by DHPLC and validation by Sanger sequencing. **(B)** *TP53* mutations, *MYC* mutations and 1q gain in 61 BL. Red = mutation, beige = wild-type, white = missing data, dark red: mutations in *MYC* residues 56-58. **(C)** Genome-wide copy number alterations in TP53wt (n=31, *left*) and TP53mut (n=25, *right*) BL. Gains are shown in green and losses are shown in red. **(D)** Detailed mirror plots show the proportion of TP53mut (red) and TP53wt (green) patients with chromosome 1q gain by genomic locus and disease. **(E)** Weighted z-scores for genes on 1q and genes not located on 1q in 4 TP53wt (green) and 4 TP53mut (red) BL cell lines. **(F)** Mean weighted z-scores of TP53wt (n=4) and TP53mut (n=4) BL cell lines from the RNAi screen with indication of genes located on chr1q and key oncogenes frequently gained in primary samples.

We next explored the profile of copy number alterations in BL (n=56, Figure 5C, S5C). Deletion of 17p13, including the *TP53* gene, was rare in BL, but co-occurred with *TP53* mutation in 5/6 cases resulting in bi-allelic p53 inactivation. Loss of the p53 activator ARF (*CDKN2A* locus on 9p21.3), that has been described as an alternative mechanism of p53 inactivation (Lindstrom et al., 2001), was rare in BL (n=1). The gain on chromosome 1q (chr1q+) affected 33.9% of BL and was the most common structural aberration besides *MYC* translocation. Unexpectedly, we observed an overrepresentation of chr1q+ in TP53wt BL (13/31, 41.9%) compared to TP53mut (6/25, 24%), suggesting a previously unappreciated connection of *TP53* status and 1q gain in BL (Figure 5B, C, S5C). A detailed analysis of the architecture of 1q CNA based on single nucleotide polymorphism (SNP) array and comparative genomic hybridization (CGH) array data showed that chromosomal gains most frequently affected the proximal part of the chromosome and an additional peak at 1q32, including the location of *MDM4* (Figure 5D). The association of chr1q gain with *TP53* mutation status was specific to BL and was not observed in DLBCL (p=1) or intermediate (p=0.654) lymphomas (Figure S5A, B).

Genes affected by the 1q gain included a number of known oncogenes, such as *BCL9*, *MCL-1*, *CKS1B* and *MDM4*, and gene clusters of the Fc receptor like (FCRL) and the Fc immunoglobulin receptor (FCGR) family that serve as potential targets for immunotherapy (Masuda et al., 2009). We therefore tested if BL cell lines from the RNAi screen were more dependent on genes on 1q (Figure 5E, D). The RNAi library covered 235 genes located on 1q including all candidate genes except for the FCRL gene family. All four TP53wt BL cell lines were previously reported to carry a 1q gain (Toujani et al., 2009). In Seraphine, the whole chromosomal arm was affected (+1q21.1qter), while partial gains were seen in BL-2 (+1q21.1q31.3), LY47 (+1q43q44), and Seraphine (+1q21.1qter). The TP53mut cell lines were diploid for 1q (Table S3). Genes on 1q were not enriched for viability genes in the group of TP53wt or TP53mut BL cell lines, respectively (Figure 5E). Notably, *MDM4* was the only gene showing *TP53*-specific viability effects after silencing (Figure 5F).

Altogether, our data support a critical role for quenching of the p53 pathway in BL preferably by mutations of *TP53* or amplification of *MDM4*, thereby identifying p53 signaling as the critical failsafe checkpoint in BL.

### *TP53* mutations and MDM4 gain inactivate the p53 pathway in primary BL

To study the functional consequences of p53 pathway aberrations, we generated a molecular signature that distinguished TP53wt and TP53mut B-cell non-Hodgkin-Lymphoma (B-NHL, n=430) using supervised hierarchical clustering (Figure 6A). The gene *CDKN2A* was significantly repressed in TP53wt BL (p<0.01), intermediate lymphoma (p<0.01) and DLBCL (p<0.01) samples (Figure 6B). Within the 50 most differentially expressed gene probes with lower expression in TP53mut patients, 28 were located on chr17p13, while 4 gene probes were located on chr1q. These findings reflect the gene dosage effect as a result of chr17p13 deletion in TP53mut and chr1q gain in TP53wt patients. Nine probes corresponding to six p53 target genes were overexpressed in TP53wt samples, demonstrating that a portion of aggressive B-NHL retain active p53 signaling. Notably, while *MDM2*, that is activated by p53 in a negative feedback loop, showed higher expression in TP53wt DLBCL (p<0.01) and BL (p=0.04) (Figure 6C), high MDM4 mRNA expression was specific to BL with TP53wt (p<0.01, Figure 6D). Expression of MDM4 in TP53wt samples was independent of chr1q+, indicating that additional mechanisms regulate MDM4 expression (Figure S6). Combined, these data provide evidence for upregulation of MDM4 in TP53wt BL as a disease driver.

**Figure 6.**
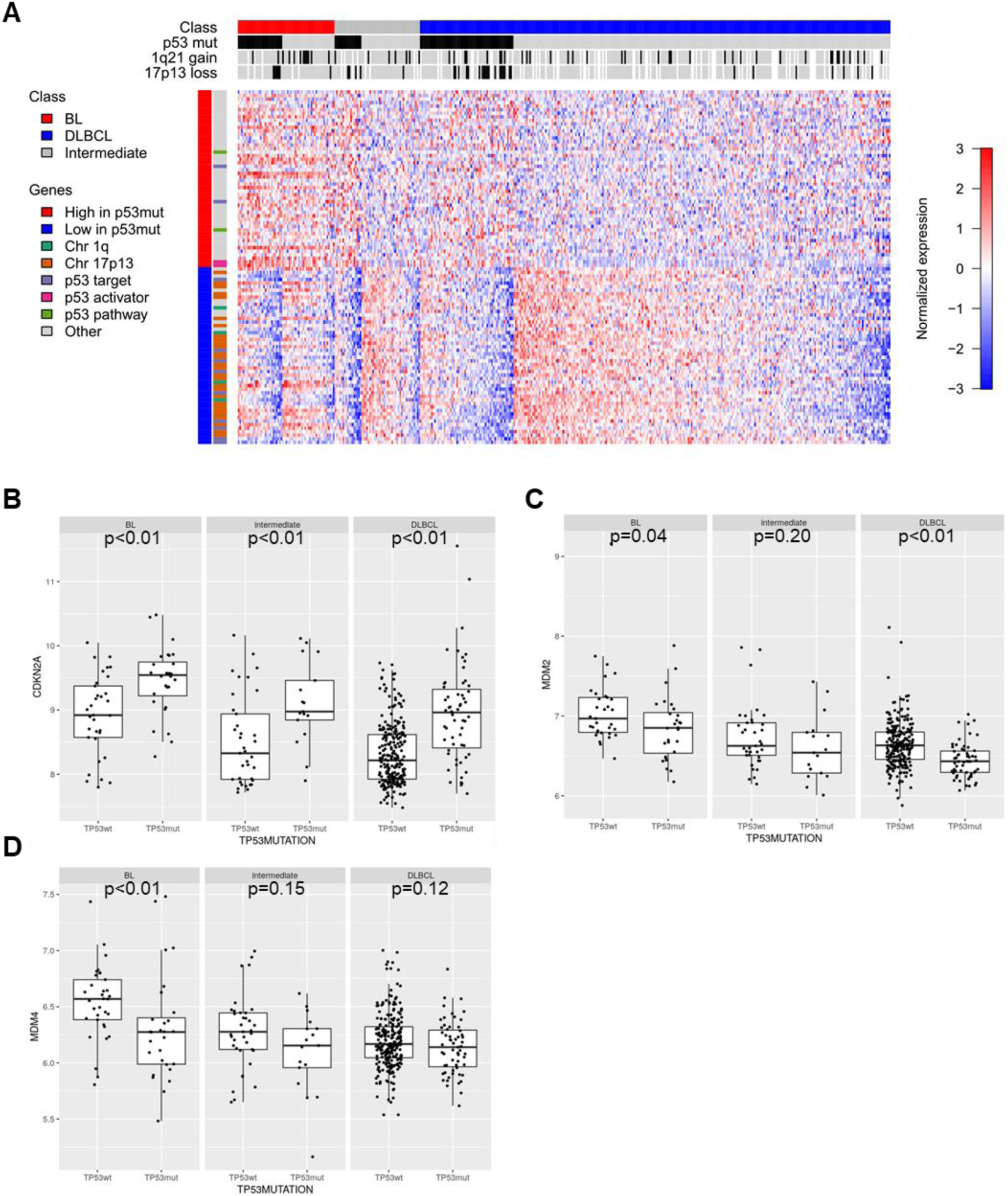
p53 pathway activation based on gene expression. **(A)** Supervised hierarchical clustering of aggressive B-NHL patients (n=412) by molecular subtype and *TP53* mutation status using the 50 gene probes with higher (red) or lower (blue) expression in TP53mut. p53 status, 17p13 deletion and 1q gain are indicated above (black = aberration, grey = normal, white = not available). **(B-D)** Differential expression of *CDKN2A* **(B)**, *MDM2* **(C)** and *MDM4* **(D)** in lymphoma subtypes stratified by *TP53* mutation status.

### *MDM4* and *TP53* mutation across cancer models

To investigate the role of chr1q gain in context of *TP53* mutations across a range of cancers, we analyzed the associations between genetic aberrations in 789 cell lines with available SNP6.0 data and *TP53* mutation data within the Cancer Cell Line Encyclopedia (Barretina et al., 2012). Chr1q32 gain was identified in 122 cell lines (15.5%) and was associated with wild-type p53 (p<0.001, 23% in TP53wt and 12% in TP53mut) (Figure 7A).

**Figure 7.**
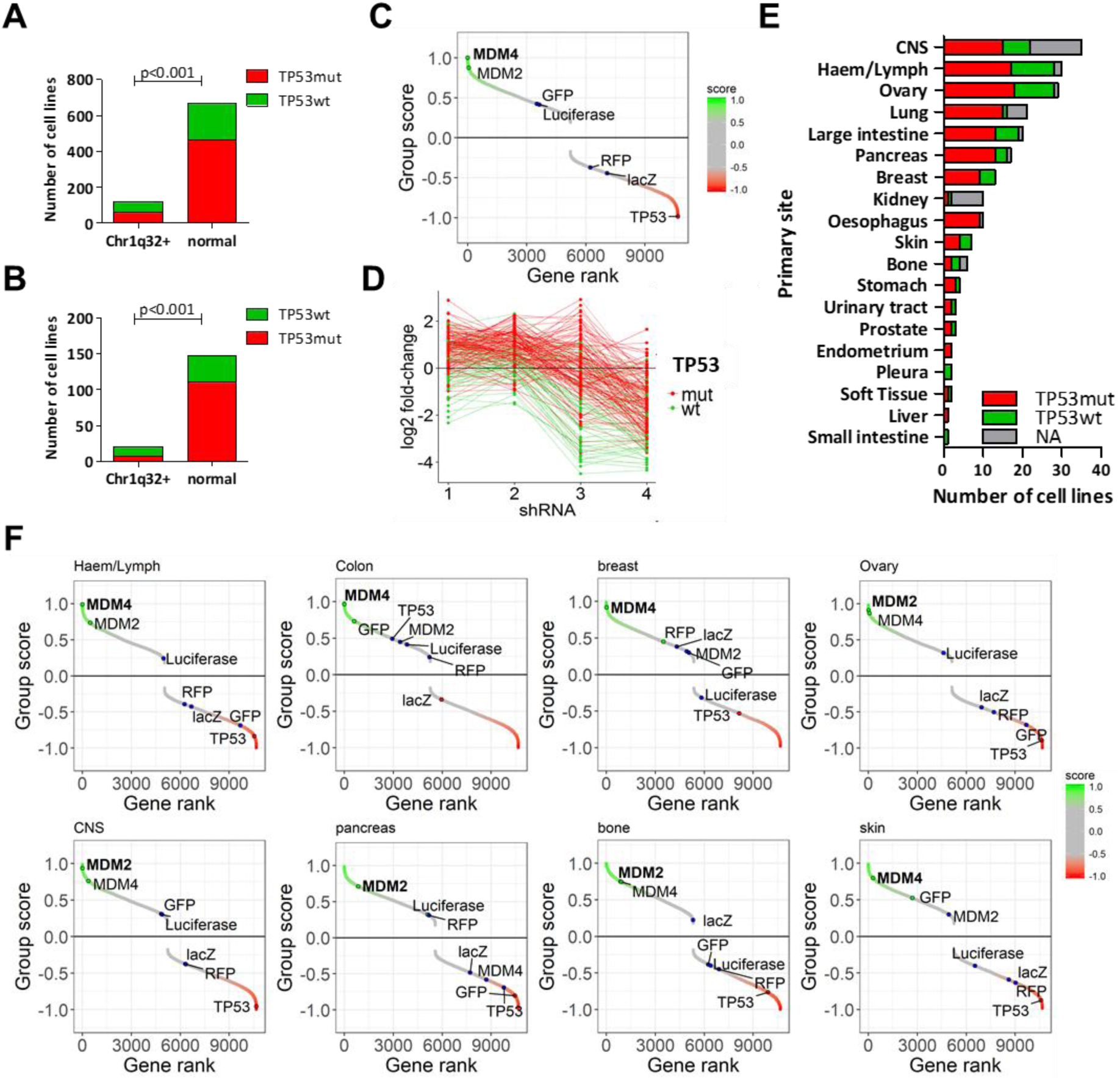
MDM4 is essential in TP53wt cancers. **(A)** Incidence of *TP53* mutation and chr1q32 gain in 789 cell lines. Information on the *TP53* status was available from COSMIC (Sanger Institute), CCLE (Broad-Novartis) and the IARC p53 data base. **(B)** Incidence of *TP53* mutation in cell lines of the Achilles project (version 2.4.3). Information on *TP53*mutation was available for 182 cell lines. **(C)** *TP53*-dependent essential genes across cancer cell lines. All genes were ranked based on their differential shRNA depletion in TP53wt (n=55) compared to TP53mut (n=127) cell lines. The genes on top of the ranking, including *MDM4* and *MDM2*, were essential in TP53wt lines. Genes that do not target human genes (GFP, RFP, luciferase and Lac-Z) serve as non-essential control genes. **(D)** Depletion of shRNAs targeting MDM4 across all cell lines. The graph shows the fold-change in shRNA expression in TP53wt (green) and TP53mut (red) cell lines. **(E)** *TP53* mutation status for 216 cell lines from the Achilles Project by cancer entity. **(F)** Entity-specific analysis of *TP53*-dependent viability genes. Gene ranking was performed for all entities that had at least two cell lines per class as described for (C).

We further combined genetic information with functional genomics data and investigated p53-dependent vulnerabilities in a set of 216 cell lines representing 19 cancer entities from the Achilles project (Cowley et al., 2014). The p53 status was available for 182 cell lines and *TP53* mutations were present in 70% of all cancer cell lines (Figure 7B, Table S5). *TP53* was identified as an essential gene in TP53mut cell lines, which is in line with its reported gain-of-function phenotype (Goldstein et al., 2011) (Figure 7C). *MDM4* was the top ranked gene leading to impaired viability of TP53wt cell lines out of more than 10,000 genes investigated (p<0.001) (Figure 7C, Table S7). All shRNAs targeting *MDM4* were strongly depleted in TP53wt cell lines (Figure 7D). Eight cancer entities were represented with at least two TP53mut and two TP53wt cell lines which allowed us to explore *MDM4* dependency in different cancer subtypes (Figure 7E, Table S7). *MDM4* was identified as an essential gene in TP53wt cell lines derived from the hematopoietic/lymphoid system (rank 1), large intestine (rank 3), breast carcinoma (rank 25) and ovarian carcinoma (rank 62) (Figure 7F). Cell lines originating from central nervous system (CNS), pancreas, bone and skin did not show p53-specific viability effects (Figure 7F).

MDM2 also showed significant shRNA depletion in TP53wt cell lines (p=0.004, rank 51, Figure 7C). Notably, p53-specific dependency on MDM2 were strongest in ovarian carcinoma (rank 20) and CNS (rank 8), suggesting disease-specific roles of MDM2 or MDM4 (Figure 7F).

Combined these data suggest a functional role for MDM4 as a critical cancer driver targeted by 1q gain across cancers.

## Discussion

Genetic studies expanded our understanding of the pathogenesis of BL and functional genomics is a powerful tool to link genetic aberrations to novel molecular targets. This study dissected context-specific essential genes in BL. We observed a strong dependence of BL on PAX5, a key B-cell transcription factor previously linked to B-cell lymphomagenesis (Cozma et al., 2007). A recent genetic perturbation screen using genome-wide CRISPR/Cas9 sgRNA libraries identified *PAX5* as one of the top candidate genes essential in two BL cell lines (Wang et al., 2015). These findings are in line with a role of PAX5 as a “lineage-survival oncogene” (Ceribelli et al., 2016; Garraway and Sellers, 2006). The increased capacity to drug transcription factors (Filippakopoulos et al., 2010) as well as the recent demonstration of the role of PAX5 as a metabolic gatekeeper (Chan et al., 2017) suggests that PAX5 targeting may provide a novel therapeutic strategy.

We systematically queried genotype-specific vulnerabilities of BL and identified oncogene dependency on TCF3 of *TCF3/ID3* mutant BL, and dependency on MYD88 and IRAK1 in a cell line with *MYD88* mutation in line with previous results in DLBCL cell lines (Ngo et al., 2011). The strongest dependency was observed for TP53wt cell lines on MDM4. While BL displays with a high rate of *TP53* mutations (higher than e.g. DLBCL), alternative mechanisms must be exploited in the cells retaining TP53wt. The dependency of TP53wt BL cell lines on MDM4 provides additional evidence for the importance of suppressing p53-mediated stress signals. Reactivation of p53 by inhibition of MDM4 has been shown to be a promising therapeutic target in melanomas (de Lange et al., 2012; Gembarska et al., 2012) and breast carcinomas (Haupt et al., 2015). We validated MDM4 as a therapeutic target in TP53wt BL in a mouse xenograft model and MDM4 silencing enhanced the cytotoxic effect of MDM2 inhibition.

Based on a pan-cancer analysis, we show that MDM4 targeting could be exploited therapeutically across cancers with TP53wt and we confirmed the association of 1q gain and cancers retaining TP53wt. Our findings provide a rational for further clinical development of dual MDM2/MDM4 inhibitors for BL and other TP53wt cancers.

Chromosome 1q gain is the most frequent copy number alteration across cancer (Beroukhim et al., 2010). Functional evidence for disease drivers targeted by 1q gain has been scarce. In BL, studies of CNA identified gains for 1q25.1 and 1q31.3 and suggested *PTPRC*, a regulator of B-cell receptor and cytokine signaling, and two annotated miRNA genes (hsa-mir-181b-1 and -213) as strong candidates (Scholtysik et al., 2010). A study of primary tumors and cell lines identified *BCA2* and *PIAS3* on 1q21.-1q21.3, *MDM4* on 1q32.1 and *AKT3* on 1q44 as possible drivers (Toujani et al., 2009). Our data provides functional evidence that 1q gain targets *MDM4* as an essential gene in TP53wt BL. Based on the incidence of *TP53* mutation and 1q gain in the disease, our findings suggest a widespread mechanism to quench p53 activity in BL, likely as a way to quench p53 mediated failsafe mechanisms elicited by MYC overexpression.

Combined, these data provide functional evidence that 1q gain targets *MDM4* as a disease driver across cancer. *TP53* mutations and 1q gains were reported as the frequent lesions associated with postmyeloproliferative-neoplasm AML and mutual exclusivity has been reported in this setting (Harutyunyan et al., 2011). Copy number changes of *MDM4* and *MDM2* have been reported to occur exclusively in bladder TP53wt tumors (Veerakumarasivam et al., 2008). A study by Monti et al identified a comprehensive set of CNAs that decreased p53 activity and perturbed cell cycle regulation in DLBCL (Monti et al., 2012).

Our data suggest that among lymphomas, BL exhibits disease specific mechanisms of p53 pathway quenching through *TP53* mutation and MDM4 overexpression. A major open question pertains to the selective advantage of MDM4 or MDM2 overexpression in TP53wt cancers. Although MDM4 and MDM2 are highly homologous, MDM4 lacks E3 ligase activity and depends on MDM2 to promote p53 degradation (Linares et al., 2003). In our study, we identified downregulation of *MYC* and upregulation of *CCND1* exclusively after MDM4 knock-down, indicating differences in pathway contribution exerted by MDM4 over MDM2. p53-independent oncogenic activities have been previously associated with MDM4. MDM4 was shown to promote pRb degradation by MDM2, with consequent E2F1 activation of the cell cycle (Zhang et al., 2015). MDM4 contributes to p53 inhibition by suppressing its transcriptional activity, and also by partnering with MDM2 to regulate p53 degradation (Francoz et al., 2006).

## Author Contribution

Conceptualization, J.H., M.S. and T.Z.;

Methodology, J.H. and M.S.;

Software, J.H., M.S., Z.H., M.Z. and O.Y.;

Validation, J.H. and M.S.;

Formal Analysis, J.H., M.S., M.R., M.K., R.K., J.L., Z.H., M.Z. and O.Y.;

Investigation, J.H., M.S., A.J., S.H., K.T., S.S., M.L. and C.P.;

Resources, J.H., M.S., M.R., A.J., R.W., S.H., K.T., Ro.Ku., J.L., S.S., Z.H., M.L., O.Y., H.H, Re.Sc., K.B., D.T.,

Ra.Kü., W.K., C.P., St.St., Ma.Lö., L.T., M.H., B.B., M.Z., Re.Si., U.K., W.H., M.K. and T.Z.;

Sample provision and profiling within the MMML, M.R., A.J., H.H., Re.Sc., Ra.Kü., W.K., C.P., St.St.,

Ma.Lö., L.T., M.H., Re.Si., M.K. and T.Z

Data Curation, J.H. and O.Y.;

Writing – Original Draft, J.H., M.S. and T.Z.;

Writing – Review & Editing, all authors;

Visualization, J.H., M.R., M.K. and S.H.;

Supervision, T.Z., U.K., B.B. and W.H.;

Funding Acquisition, T.Z.

## Acknowledgements

The work was supported by the *Helmholtz Virtual Institute* „Understanding and overcoming resistance to apoptosis and therapy in leukemia”, the Helmholtz initiative *iMed* on Personalized Medicine, the European Union (FP7 projects *Radiant, Systems Microscopy*, Horizon 2020 project *SOUND*) and the “Monique Dornonville de la Cour – Stiftung“.

The “Deutsche Krebshilfe” supported TZ (“Mildred-Scheel” Professorship), ML (“Mildred-Scheel” Fellowship) and the “Molecular Mechanisms of Malignant Lymphoma – MMML” consortium. RS/RW received infrastructural support by the “KinderKrebsInitiative Buchholz Holm-Seppensen”.

For technical support and expertise we thank the DKFZ Genomics and Proteomics Core Facility. We thank Hanno Glimm, Stefan Fröhling, Daniela Richter, Roland Eils, Peter Lichter, Stephan Wolf, Katja Beck and Janna Kirchhof for infrastructure and program development within DKFZ-HIPO and NCT POP, and Tina Uhrig for technical assistance and Agnes Hotz-Wagenblatt for shRNA alignment. We thank Anna Jauch for FISH analysis in BL cell lines. We thank Henry-Jacques Delecluse and Astrid Hofmann for staining of EBV proteins in BL cell lines to determine the EBV status and latency phase.

---

**Supplementary Figure 1.** Extended shRNA screening results.

**Supplementary Figure 2.** Fold-changes of shRNAs targeting proteasomal subunits in BL cell lines.

**Supplementary Figure 3.** p53 impairment in Seraphine-p53ko cell line.

**Supplementary Figure 4.** Differential gene expression between MDM2 and MDM4 knock-down in Seraphine-p53wt.

**Supplementary Figure 5.** Copy number alterations in primary B-NHL samples.

**Supplementary Figure 6.** Basal mRNA expression in primary B-NHL samples stratified by p53 mutation status and 1q gain.

**Supplementary Table S1.** List of cell lines screened by Cellecta Inc..

**Supplementary Table S2.** Gene scores from class comparison.

**Supplementary Table S3.** Mutations in BL cell lines.

**Supplementary Table S4.** Gene expression profiling after MDM4 and MDM2 knock-down in Seraphine-TP53wt and Seraphine-p53ko.

**Supplementary Table S5.** Annotation of patient cohort.

**Supplementary Table S6.** Achilles cell lines with *TP53* and chr1q annotation.

**Supplementary Table S7.** Achilles cell lines gene scores.

## Methods

### Cell culture

BJAB, BL-2, CA46, Namalwa, Ramos, Raji, BL-41, DogKit, DG-75 and Gumbus were obtained from DSMZ (Braunschweig, Germany), BL7, BL60, LY47 were provided by G.M. Lenoir (IARC, Lyon, France), Salina, Seraphine, and Cheptanges were provided by A. Rickinson, (Birmingham, UK) and 293T/17 by Stefan Fröhling (DKFZ, Heidelberg, Germany). All cell lines were maintained under standard conditions. Cell line authentification was performed using Multiplex Cell Authentification by Multiplexion (Heidelberg, Germany) as described previously (Castro et al., 2013). The SNP profiles matched with known ones in the databse or were unique. Cell lines were tested for contamination using the Multiplexion cell Contamination test (Heidelberg, Germany) as described previously (Schmitt and Pawlita, 2009).

### RNAi screen

The RNAi screen was performed as described previously using the Cellecta human Module I pooled lentiviral shRNA library (Slabicki et al., 2016). Cells were harvested on day 2 (one replicate) or day 14 (2 replicates) post-transduction. p-values for shRNA depletion were calculated with the edgeR package (Dai et al., 2014) and shRNA p-values were collapsed into gene scores using weighted Z-transformation (Kim and Tan, 2012). For comparison of differential shRNA effects between two groups, log2-transformed shRNA fold-changes were scaled with peak median absolute deviation (PMAD) normalization using the GenePattern module NormLines (Cheung et al., 2011; Reich et al., 2006). Using the GENE-E software (https://software.broadinstitute.org/GENE-E/index.html), p-values for differential shRNA effects were calculated and collapsed into gene scores using Kolmogorov-Smirnov statistics (Cheung et al., 2011). RNAi results in non-BL cell lines were provided by Cellecta Inc. as raw read counts. Genome-wide RNAi results in 216 cell lines were provided as log2-transformed shRNA fold-changes (Cowley et al., 2014). Genetic information on cell lines was extracted from CCLE (https://portals.broadinstitute.org/ccle/home) and COSMIC (GDSC, http://www.cancerrxgene.org/). shRNAs were mapped against the human transcriptome (Ensembl v75) using Tophat2 algorithm and genes that scored with shRNAs targeting non-overlapping sequences were considered as candidate genes. For single gene knock-downs, shRNAs were expressed in the pRSI12-U6-(sh)-UbiC-TagRFP-2A-Puro vector backbone. Packaging of constructs into lentiviral particles was performed as described previously (Slabicki et al., 2016). shMDM4_1: 5’-GTTCACTGTTAAAGAGGTCAT-3’; shMDM4_2: 5’-CACCTAGAAGTAATGGCTCAA-3’; shMDM2_1: 5’-CTTTGGTAGTGGAATAGTGAA-3’.

### RNA sequencing

mRNA sequencing libraries were prepared from 1 µg of total RNA using Illumina TruSeq R RNA sample preparation v2 (Illumina, San Diego, CA, USA) and multiplexed samples were sequenced on Illumina HiSeq 2000 by the Genomics and Proteomics Core Facility (DKFZ). Sequences were mapped against the human reference genome hg19 using the STAR alignment tool. HT-Seq count was used to calculate gene-level counts and RPKM (RPKM = (10^9 * C)/(N * L); C=Number of reads mapped to a gene; N=Total Mapped Reads on exon; L=Exon length in base-pairs for a gene) with annotation GENCODE version 19 GTF. Mutations were called from RNA sequencing and targeted resequencing as described previously (Hullein et al., 2013; International Cancer Genome Consortium PedBrain Tumor, 2016).

### qRT-PCR

Total RNA was isolated using the RNeasy Mini Kit (Qiagen) with on-column DNase I (Qiagen) digestion and 500 ng of total RNA was subsequently reverse-transcribed by Super-Script III First-Strand Synthesis Supermix (Invitrogen). QuantiFast SYBR Green RT-PCR (Qiagen) or Power SYBR Green Master Mix (Applied Biosystems) was used for quantitative PCR reaction on a LightCycler 480 Real-Time PCR System (Roche Applied Sciences) and analyzed using LightCycler 480 Software, Version 1.5 (Roche). qRT-PCR primers were MDM4_fwd: 5’-TGAAAGACCCAAGCCCTCT-3’; MDM4_rev: 5’-CGAGAGTCTGAGCAGCATCTG-3’; TP53_fwd: 5’-GGAGCACTAAGCGAGCACTG-3’; TP53_rev: 5’-CACGGATCTGAAGGGTGAAA-3’; MDM2_fwd: 5’-CAGTAGCAGTGAATCTACAGGGA-3’; MDM2_rev: 5’-CTGATCCAACCAATCACCTGAAT-3’; GAPDH_fwd: 5’-ACCCAGAAGACTGTGGATGG-3’; GAPDH_rev: 5’-TCTAGACGGCAGGTCAGGTC-3’.

### Immunoblot analysis

Following antibodies were used in this study for immunoblot analysis: anti-MDM4 (cat. 04-1555, Merck-Millipore Billerica); anti-MDM2 (cat. OP46, Merck-Millipore), anti-GAPDH (cat. ab9485, Abcam), anti-p53 (cat. 554294, BD Pharmingen), anti-p21 (cat. 556431, Santa Cruz Biotechnology), anti-cleaved PARP (cat. 9546, Cell Signaling), anti-mouse IgG DyLight800 (cat. 5257, Cell Signaling), anti-rabbit IgG (H+L) DyLight680 (cat. 5366, Cell Signaling) and the LI-COR Odyssey Infrared Imaging System (Cell Signaling) was used for detection.

### Generation of isogenic cell lines

sgRNAs targeting p53 were cloned into lentiCRISPRv2, which was a gift from Feng Zhang (Addgene, Cambridge, MA, USA, plasmid #52961) (Sanjana et al., 2014). Seraphine cells modified with lentiCRISPRv2-sgTP53 were selected using puromycin and Nutlin-3. sgTP53: 5’-CCCCTTGCCGTCCCAAGCAA-3’.

### p53 staining

Cells were fixed with the FIX & PERM Cell Fixation and Cell Permeabilization Kit (Thermo Fisher Scientific) according to the manufacture’s protocol. Subsequently, cells were stained with anti-p53 antibody (cat. 554293, BD Pharmingen) and subjected to flow cytometry.

### Cell cycle profiling/apoptosis assay

Analysis of cell cycle phases was performed using the BrdU Flow Kit (BD Pharmingen), including Annexin-V to stain for apoptotic cells according to the manufacturer’s protocols.

### Gene expression analysis

Global gene expression changes after MDM4 and MDM2 knock-down were analyzed by hybridization of total RNA on a Illumina BeadChip HumanHT-12-v4 containing >47,000 probes for 31,000 annotated human genes. To identify biological processes correlated with differentially expressed genes, the Gene Set Enrichment Analysis (GSEA) software was used with the C2 and H gene sets from the MSigDB database (http://software.broadinstitute.org/gsea/msigdb) (Mootha et al., 2003; Subramanian et al., 2005).

### Xenograft model

Animal studies were performed in agreement with the Guide for Care and Use of Laboratory Animals published by the US National Institutes of Health (NIH Publication n. 85–23, revised 1996), in compliance with the German law on the protection of animals, and with the approval of the regional authorities responsible (Regierung von Oberbayern). The *in-vivo* experiments were performed as published previously (Herhaus et al., 2016). Briefly, Seraphine-TP53wt, Seraphine-p53ko and Raji cell lines were infected *in vitro* with shNT or shMDM4 aiming at > 80% transduction efficiency. 1×10^7 cells were subcutaneously injected into flanks of immunodeficient mice. Tumor growth was monitored by FDG-PET after 11 or 16 days depending on the graft efficiency and mice were sacrificed.

### Luminescent growth assay

Cell viability and growth were measured by detection of ATP level using the CellTiter-Glo luminescent assay (Promega, Madison, WI) as described (Dietrich et al., 2018).

### Primary sample analysis

Copy number alterations were analyzed by CGH using a BAC/PAC array that consisted of 2799 DNA fragments as described elsewhere (Fiegler et al., 2003; Schwaenen et al., 2004). Interphase FISH analysis was performed on paraffin-embedded or frozen tissue sections to determine *MYC*, *BCL2* and *BCL6* translocations to IG regions. *TP53* mutations were determined by DHPLC and sequencing of exons 4-10 of the coding region (Zenz et al., 2010). The expression data of primary samples was downloaded from the Gene Expression Omnibus (http://www.ncbi.nlm.nih.gov/geo, accession numbers GSE43677, GSE21597). Patients were classified into BL, DLBCL and an intermediate group based on a previously described molecular signature (Hummel et al., 2006). For all samples, tumor cell content exceeded 70%.

## References

Baldi, A., Piccolo, M. T., Boccellino, M. R., Donizetti, A., Cardillo, I., La Porta, R., Quagliuolo, L., Spugnini, E. P., Cordero, F., Citro, G., et al. (2011). Apoptosis induced by piroxicam plus cisplatin combined treatment is triggered by p21 in mesothelioma. PloS one 6, e23569.

Barretina, J., Caponigro, G., Stransky, N., Venkatesan, K., Margolin, A. A., Kim, S., Wilson, C. J., Lehar, J., Kryukov, G. V., Sonkin, D., et al. (2012). The Cancer Cell Line Encyclopedia enables predictive modelling of anticancer drug sensitivity. Nature 483, 603–607.

Bartek, J., Bartkova, J., Vojtesek, B., Staskova, Z., Lukas, J., Rejthar, A., Kovarik, J., Midgley, C. A., Gannon, J. V., and Lane, D. P. (1991). Aberrant expression of the p53 oncoprotein is a common feature of a wide spectrum of human malignancies. Oncogene 6, 1699–1703.

Beroukhim, R., Mermel, C. H., Porter, D., Wei, G., Raychaudhuri, S., Donovan, J., Barretina, J., Boehm, J. S., Dobson, J., Urashima, M., et al. (2010). The landscape of somatic copy-number alteration across human cancers. Nature 463, 899–905.

Bhatia, K. G., Gutierrez, M. I., Huppi, K., Siwarski, D., and Magrath, I. T. (1992). The pattern of p53 mutations in Burkitt’s lymphoma differs from that of solid tumors. Cancer Res 52, 4273–4276.

Blagosklonny, M. V., Toretsky, J., Bohen, S., and Neckers, L. (1996). Mutant conformation of p53 translated in vitro or in vivo requires functional HSP90. Proceedings of the National Academy of Sciences of the United States of America 93, 8379–8383.

Burgess, A., Chia, K. M., Haupt, S., Thomas, D., Haupt, Y., and Lim, E. (2016). Clinical Overview of MDM2/X-Targeted Therapies. Frontiers in oncology 6, 7.

Castro, F., Dirks, W. G., Fahnrich, S., Hotz-Wagenblatt, A., Pawlita, M., and Schmitt, M. (2013). High-throughput SNP-based authentication of human cell lines. Int J Cancer 132, 308–314.

Ceribelli, M., Hou, Z. E., Kelly, P. N., Huang, D. W., Wright, G., Ganapathi, K., Evbuomwan, M. O., Pittaluga, S., Shaffer, A. L., Marcucci, G., et al. (2016). A Druggable TCF4- and BRD4-Dependent Transcriptional Network Sustains Malignancy in Blastic Plasmacytoid Dendritic Cell Neoplasm. Cancer cell 30, 764–778.

Chan, L. N., Chen, Z., Braas, D., Lee, J. W., Xiao, G., Geng, H., Cosgun, K. N., Hurtz, C., Shojaee, S., Cazzaniga, V., et al. (2017). Metabolic gatekeeper function of B-lymphoid transcription factors. Nature 542, 479–483.

Cheung, H. W., Cowley, G. S., Weir, B. A., Boehm, J. S., Rusin, S., Scott, J. A., East, A., Ali, L. D., Lizotte, P. H., Wong, T. C., et al. (2011). Systematic investigation of genetic vulnerabilities across cancer cell lines reveals lineage-specific dependencies in ovarian cancer. Proceedings of the National Academy of Sciences of the United States of America 108, 12372–12377.

Cowley, G. S., Weir, B. A., Vazquez, F., Tamayo, P., Scott, J. A., Rusin, S., East-Seletsky, A., Ali, L. D., Gerath, W. F., Pantel, S. E., et al. (2014). Parallel genome-scale loss of function screens in 216 cancer cell lines for the identification of context-specific genetic dependencies. Scientific data 1, 140035.

Cozma, D., Yu, D., Hodawadekar, S., Azvolinsky, A., Grande, S., Tobias, J. W., Metzgar, M. H., Paterson, J., Erikson, J., Marafioti, T., et al. (2007). B cell activator PAX5 promotes lymphomagenesis through stimulation of B cell receptor signaling. The Journal of clinical investigation 117, 2602–2610.

Dai, Z., Sheridan, J. M., Gearing, L. J., Moore, D. L., Su, S., Dickins, R. A., Blewitt, M. E., and Ritchie, M. E. (2014). shRNA-seq data analysis with edgeR. F1000Res 3, 95.

Dalla-Favera, R., Bregni, M., Erikson, J., Patterson, D., Gallo, R. C., and Croce, C. M. (1982). Human c-myc onc gene is located on the region of chromosome 8 that is translocated in Burkitt lymphoma cells. Proceedings of the National Academy of Sciences of the United States of America 79, 7824–7827.

de Lange, J., Teunisse, A. F., Vries, M. V., Lodder, K., Lam, S., Luyten, G. P., Bernal, F., Jager, M. J., and Jochemsen, A. G. (2012). High levels of Hdmx promote cell growth in a subset of uveal melanomas. Am J Cancer Res 2, 492–507.

Dietrich, S., Oles, M., Lu, J., Sellner, L., Anders, S., Velten, B., Wu, B., Hullein, J., da Silva Liberio, M., Walther, T., et al. (2018). Drug-perturbation-based stratification of blood cancer. J Clin Invest 128, 427–445.

Evan, G. I., Wyllie, A. H., Gilbert, C. S., Littlewood, T. D., Land, H., Brooks, M., Waters, C. M., Penn, L. Z., and Hancock, D. C. (1992). Induction of apoptosis in fibroblasts by c-myc protein. Cell 69, 119–128.

Farrell, P. J., Allan, G. J., Shanahan, F., Vousden, K. H., and Crook, T. (1991). p53 is frequently mutated in Burkitt’s lymphoma cell lines. The EMBO journal 10, 2879–2887.

Fiegler, H., Carr, P., Douglas, E. J., Burford, D. C., Hunt, S., Scott, C. E., Smith, J., Vetrie, D., Gorman, P., Tomlinson, I. P., and Carter, N. P. (2003). DNA microarrays for comparative genomic hybridization based on DOP-PCR amplification of BAC and PAC clones. Genes, chromosomes & cancer 36, 361–374.

Filippakopoulos, P., Qi, J., Picaud, S., Shen, Y., Smith, W. B., Fedorov, O., Morse, E. M., Keates, T., Hickman, T. T., Felletar, I., et al. (2010). Selective inhibition of BET bromodomains. Nature 468, 1067–1073.

Fish, K., Sora, R. P., Schaller, S. J., Longnecker, R., and Ikeda, M. (2017). EBV latent membrane protein 2A orchestrates p27(kip1) degradation via Cks1 to accelerate MYC-driven lymphoma in mice. Blood 130, 2516–2526.

Gaidano, G., Ballerini, P., Gong, J. Z., Inghirami, G., Neri, A., Newcomb, E. W., Magrath, I. T., Knowles, D. M., and Dalla-Favera, R. (1991). p53 mutations in human lymphoid malignancies: association with Burkitt lymphoma and chronic lymphocytic leukemia. Proceedings of the National Academy of Sciences of the United States of America 88, 5413–5417.

Garraway, L. A., and Sellers, W. R. (2006). Lineage dependency and lineage-survival oncogenes in human cancer. Nature reviews Cancer 6, 593–602.

Gembarska, A., Luciani, F., Fedele, C., Russell, E. A., Dewaele, M., Villar, S., Zwolinska, A., Haupt, S., de Lange, J., Yip, D., et al. (2012). MDM4 is a key therapeutic target in cutaneous melanoma. Nature medicine.

Goldstein, I., Marcel, V., Olivier, M., Oren, M., Rotter, V., and Hainaut, P. (2011). Understanding wild-type and mutant p53 activities in human cancer: new landmarks on the way to targeted therapies. Cancer Gene Ther 18, 2–11.

Graves, B., Thompson, T., Xia, M., Janson, C., Lukacs, C., Deo, D., Di Lello, P., Fry, D., Garvie, C., Huang, K. S., et al. (2012). Activation of the p53 pathway by small-molecule-induced MDM2 and MDMX dimerization. Proceedings of the National Academy of Sciences of the United States of America 109, 11788–11793.

Halasi, M., Pandit, B., and Gartel, A. L. (2014). Proteasome inhibitors suppress the protein expression of mutant p53. Cell cycle 13, 3202–3206.

Hart, T., Brown, K. R., Sircoulomb, F., Rottapel, R., and Moffat, J. (2014). Measuring error rates in genomic perturbation screens: gold standards for human functional genomics. Molecular systems biology 10, 733.

Harutyunyan, A., Klampfl, T., Cazzola, M., and Kralovics, R. (2011). p53 lesions in leukemic transformation. N Engl J Med 364, 488–490.

Haupt, S., Buckley, D., Pang, J. M., Panimaya, J., Paul, P. J., Gamell, C., Takano, E. A., Ying Lee, Y., Hiddingh, S., Rogers, T. M., et al. (2015). Targeting Mdmx to treat breast cancers with wild-type p53. Cell death & disease 6, e1821.

Helming, K. C., Wang, X., Wilson, B. G., Vazquez, F., Haswell, J. R., Manchester, H. E., Kim, Y., Kryukov, G. V., Ghandi, M., Aguirre, A. J., et al. (2014). ARID1B is a specific vulnerability in ARID1A-mutant cancers. Nature medicine 20, 251–254.

Hemann, M. T., Bric, A., Teruya-Feldstein, J., Herbst, A., Nilsson, J. A., Cordon-Cardo, C., Cleveland, J. L., Tansey, W. P., and Lowe, S. W. (2005). Evasion of the p53 tumour surveillance network by tumour-derived MYC mutants. Nature 436, 807–811.

Heminger, K., Markey, M., Mpagi, M., and Berberich, S. J. (2009). Alterations in gene expression and sensitivity to genotoxic stress following HdmX or Hdm2 knockdown in human tumor cells harboring wild-type p53. Aging (Albany NY) 1, 89–108.

Herhaus, P., Habringer, S., Philipp-Abbrederis, K., Vag, T., Gerngross, C., Schottelius, M., Slotta-Huspenina, J., Steiger, K., Altmann, T., Weisser, T., et al. (2016). Targeted positron emission tomography imaging of CXCR4 expression in patients with acute myeloid leukemia. Haematologica 101, 932–940.

Hu, B., Gilkes, D. M., Farooqi, B., Sebti, S. M., and Chen, J. (2006). MDMX overexpression prevents p53 activation by the MDM2 inhibitor Nutlin. The Journal of biological chemistry 281, 33030–33035.

Hullein, J., Jethwa, A., Stolz, T., Blume, C., Sellner, L., Sill, M., Langer, C., Jauch, A., Paruzynski, A., von Kalle, C., et al. (2013). Next-generation sequencing of cancer consensus genes in lymphoma. Leuk Lymphoma 54, 1831–1835.

Hummel, M., Bentink, S., Berger, H., Klapper, W., Wessendorf, S., Barth, T. F., Bernd, H. W., Cogliatti, S. B., Dierlamm, J., Feller, A. C., et al. (2006). A biologic definition of Burkitt’s lymphoma from transcriptional and genomic profiling. N Engl J Med 354, 2419–2430.

International Cancer Genome Consortium PedBrain Tumor, P. (2016). Recurrent MET fusion genes represent a drug target in pediatric glioblastoma. Nature medicine 22, 1314–1320.

Kim, J., and Tan, A. C. (2012). BiNGS!SL-seq: A Bioinformatics Pipeline for the Analysis and Interpretation of Deep Sequencing Genome-Wide Synthetic Lethal Screen. Next Generation Microarray Bioinformatics: Methods and Protocols 802, 389–398.

Kretzmer, H., Bernhart, S. H., Wang, W., Haake, A., Weniger, M. A., Bergmann, A. K., Betts, M. J., Carrillo-de-Santa-Pau, E., Doose, G., Gutwein, J., et al. (2015). DNA methylome analysis in Burkitt and follicular lymphomas identifies differentially methylated regions linked to somatic mutation and transcriptional control. Nature genetics 47, 1316–1325.

Li, D., Marchenko, N. D., Schulz, R., Fischer, V., Velasco-Hernandez, T., Talos, F., and Moll, U. M. (2011). Functional inactivation of endogenous MDM2 and CHIP by HSP90 causes aberrant stabilization of mutant p53 in human cancer cells. Mol Cancer Res 9, 577–588.

Linares, L. K., Hengstermann, A., Ciechanover, A., Muller, S., and Scheffner, M. (2003). HdmX stimulates Hdm2-mediated ubiquitination and degradation of p53. Proceedings of the National Academy of Sciences of the United States of America 100, 12009–12014.

Lindstrom, M. S., Klangby, U., and Wiman, K. G. (2001). p14ARF homozygous deletion or MDM2 overexpression in Burkitt lymphoma lines carrying wild type p53. Oncogene 20, 2171–2177.

Love, C., Sun, Z., Jima, D., Li, G., Zhang, J., Miles, R., Richards, K. L., Dunphy, C. H., Choi, W. W., Srivastava, G., et al. (2012). The genetic landscape of mutations in Burkitt lymphoma. Nature genetics.

Marcotte, R., Sayad, A., Brown, K. R., Sanchez-Garcia, F., Reimand, J., Haider, M., Virtanen, C., Bradner, J. E., Bader, G. D., Mills, G. B., et al. (2016). Functional Genomic Landscape of Human Breast Cancer Drivers, Vulnerabilities, and Resistance. Cell 164, 293–309.

Masuda, A., Yoshida, M., Shiomi, H., Morita, Y., Kutsumi, H., Inokuchi, H., Mizuno, S., Nakamura, A., Takai, T., Blumberg, R. S., and Azuma, T. (2009). Role of Fc Receptors as a therapeutic target. Inflammation & allergy drug targets 8, 80–86.

Meyer, N., Kim, S. S., and Penn, L. Z. (2006). The Oscar-worthy role of Myc in apoptosis. Semin Cancer Biol 16, 275–287.

Mohr, S. E., Smith, J. A., Shamu, C. E., Neumuller, R. A., and Perrimon, N. (2014). RNAi screening comes of age: improved techniques and complementary approaches. Nature reviews Molecular cell biology 15, 591–600.

Molyneux, E. M., Rochford, R., Griffin, B., Newton, R., Jackson, G., Menon, G., Harrison, C. J., Israels, T., and Bailey, S. (2012). Burkitt’s lymphoma. Lancet 379, 1234–1244.

Monti, S., Chapuy, B., Takeyama, K., Rodig, S. J., Hao, Y., Yeda, K. T., Inguilizian, H., Mermel, C., Currie, T., Dogan, A., et al. (2012). Integrative analysis reveals an outcome-associated and targetable pattern of p53 and cell cycle deregulation in diffuse large B cell lymphoma. Cancer cell 22, 359–372.

Mootha, V. K., Lindgren, C. M., Eriksson, K. F., Subramanian, A., Sihag, S., Lehar, J., Puigserver, P., Carlsson, E., Ridderstrale, M., Laurila, E., et al. (2003). PGC-1alpha-responsive genes involved in oxidative phosphorylation are coordinately downregulated in human diabetes. Nature genetics 34, 267–273.

Muller, P., Hrstka, R., Coomber, D., Lane, D. P., and Vojtesek, B. (2008). Chaperone-dependent stabilization and degradation of p53 mutants. Oncogene 27, 3371–3383.

Ngo, V. N., Young, R. M., Schmitz, R., Jhavar, S., Xiao, W., Lim, K. H., Kohlhammer, H., Xu, W., Yang, Y., Zhao, H., et al. (2011). Oncogenically active MYD88 mutations in human lymphoma. Nature 470, 115–119.

O’Connor, P. M., Jackman, J., Jondle, D., Bhatia, K., Magrath, I., and Kohn, K. W. (1993). Role of the p53 tumor suppressor gene in cell cycle arrest and radiosensitivity of Burkitt’s lymphoma cell lines. Cancer Res 53, 4776–4780.

Patton, J. T., Mayo, L. D., Singhi, A. D., Gudkov, A. V., Stark, G. R., and Jackson, M. W. (2006). Levels of HdmX expression dictate the sensitivity of normal and transformed cells to Nutlin-3. Cancer Res 66, 3169–3176.

Reich, M., Liefeld, T., Gould, J., Lerner, J., Tamayo, P., and Mesirov, J. P. (2006). GenePattern 2.0. Nature genetics 38, 500–501.

Richter, J., Schlesner, M., Hoffmann, S., Kreuz, M., Leich, E., Burkhardt, B., Rosolowski, M., Ammerpohl, O., Wagener, R., Bernhart, S. H., et al. (2012). Recurrent mutation of the ID3 gene in Burkitt lymphoma identified by integrated genome, exome and transcriptome sequencing. Nature genetics.

Saha, A., Murakami, M., Kumar, P., Bajaj, B., Sims, K., and Robertson, E. S. (2009). Epstein-Barr virus nuclear antigen 3C augments Mdm2-mediated p53 ubiquitination and degradation by deubiquitinating Mdm2. Journal of virology 83, 4652–4669.

Salaverria, I., Zettl, A., Bea, S., Hartmann, E. M., Dave, S. S., Wright, G. W., Boerma, E. J., Kluin, P. M., Ott, G., Chan, W. C., et al. (2008). Chromosomal alterations detected by comparative genomic hybridization in subgroups of gene expression-defined Burkitt’s lymphoma. Haematologica 93, 1327–1334.

Sanjana, N. E., Shalem, O., and Zhang, F. (2014). Improved vectors and genome-wide libraries for CRISPR screening. Nat Methods 11, 783–784.

Schmitt, M., and Pawlita, M. (2009). High-throughput detection and multiplex identification of cell contaminations. Nucleic acids research 37, e119.

Schmitz, R., Young, R. M., Ceribelli, M., Jhavar, S., Xiao, W., Zhang, M., Wright, G., Shaffer, A. L., Hodson, D. J., Buras, E., et al. (2012). Burkitt lymphoma pathogenesis and therapeutic targets from structural and functional genomics. Nature 490, 116–120.

Scholtysik, R., Kreuz, M., Klapper, W., Burkhardt, B., Feller, A. C., Hummel, M., Loeffler, M., Rosolowski, M., Schwaenen, C., Spang, R., et al. (2010). Detection of genomic aberrations in molecularly defined Burkitt’s lymphoma by array-based, high resolution, single nucleotide polymorphism analysis. Haematologica 95, 2047–2055.

Schwaenen, C., Nessling, M., Wessendorf, S., Salvi, T., Wrobel, G., Radlwimmer, B., Kestler, H. A., Haslinger, C., Stilgenbauer, S., Dohner, H., et al. (2004). Automated array-based genomic profiling in chronic lymphocytic leukemia: development of a clinical tool and discovery of recurrent genomic alterations. Proceedings of the National Academy of Sciences of the United States of America 101, 1039–1044.

Shvarts, A., Bazuine, M., Dekker, P., Ramos, Y. F., Steegenga, W. T., Merckx, G., van Ham, R. C., van der Houven van Oordt, W., van der Eb, A. J., and Jochemsen, A. G. (1997). Isolation and identification of the human homolog of a new p53-binding protein, Mdmx. Genomics 43, 34–42.

Slabicki, M., Lee, K. S., Jethwa, A., Sellner, L., Sacco, F., Walther, T., Hullein, J., Dietrich, S., Wu, B., Lipka, D. B., et al. (2016). Dissection of CD20 regulation in lymphoma using RNAi. Leukemia 30, 2409–2412.

Subramanian, A., Tamayo, P., Mootha, V. K., Mukherjee, S., Ebert, B. L., Gillette, M. A., Paulovich, A., Pomeroy, S. L., Golub, T. R., Lander, E. S., and Mesirov, J. P. (2005). Gene set enrichment analysis: a knowledge-based approach for interpreting genome-wide expression profiles. Proceedings of the National Academy of Sciences of the United States of America 102, 15545–15550.

Taub, R., Kirsch, I., Morton, C., Lenoir, G., Swan, D., Tronick, S., Aaronson, S., and Leder, P. (1982). Translocation of the c-myc gene into the immunoglobulin heavy chain locus in human Burkitt lymphoma and murine plasmacytoma cells. Proceedings of the National Academy of Sciences of the United States of America 79, 7837–7841.

Toujani, S., Dessen, P., Ithzar, N., Danglot, G., Richon, C., Vassetzky, Y., Robert, T., Lazar, V., Bosq, J., Da Costa, L., et al. (2009). High resolution genome-wide analysis of chromosomal alterations in Burkitt’s lymphoma. PloS one 4, e7089.

Tsherniak, A., Vazquez, F., Montgomery, P. G., Weir, B. A., Kryukov, G., Cowley, G. S., Gill, S., Harrington, W. F., Pantel, S., Krill-Burger, J. M., et al. (2017). Defining a Cancer Dependency Map. Cell 170, 564–576 e516.

Vassilev, L. T., Vu, B. T., Graves, B., Carvajal, D., Podlaski, F., Filipovic, Z., Kong, N., Kammlott, U., Lukacs, C., Klein, C., et al. (2004). In vivo activation of the p53 pathway by small-molecule antagonists of MDM2. Science 303, 844–848.

Veerakumarasivam, A., Scott, H. E., Chin, S. F., Warren, A., Wallard, M. J., Grimmer, D., Ichimura, K., Caldas, C., Collins, V. P., Neal, D. E., and Kelly, J. D. (2008). High-resolution array-based comparative genomic hybridization of bladder cancers identifies mouse double minute 4 (MDM4) as an amplification target exclusive of MDM2 and TP53. Clinical cancer research : an official journal of the American Association for Cancer Research 14, 2527–2534.

Vijayakumaran, R., Tan, K. H., Miranda, P. J., Haupt, S., and Haupt, Y. (2015). Regulation of Mutant p53 Protein Expression. Frontiers in oncology 5, 284.

Walerych, D., Lisek, K., Sommaggio, R., Piazza, S., Ciani, Y., Dalla, E., Rajkowska, K., Gaweda-Walerych, K., Ingallina, E., Tonelli, C., et al. (2016). Proteasome machinery is instrumental in a common gain-of-function program of the p53 missense mutants in cancer. Nature cell biology 18, 897–909.

Wang, T., Birsoy, K., Hughes, N. W., Krupczak, K. M., Post, Y., Wei, J. J., Lander, E. S., and Sabatini, D. M. (2015). Identification and characterization of essential genes in the human genome. Science.

Zenz, T., Eichhorst, B., Busch, R., Denzel, T., Habe, S., Winkler, D., Buhler, A., Edelmann, J., Bergmann, M., Hopfinger, G., et al. (2010). TP53 mutation and survival in chronic lymphocytic leukemia. Journal of clinical oncology : official journal of the American Society of Clinical Oncology 28, 4473–4479.

Zhang, X. K., Moussa, O., LaRue, A., Bradshaw, S., Molano, I., Spyropoulos, D. D., Gilkeson, G. S., and Watson, D. K. (2008). The transcription factor Fli-1 modulates marginal zone and follicular B cell development in mice. Journal of immunology 181, 1644–1654.

Zochodne, B., Truong, A. H., Stetler, K., Higgins, R. R., Howard, J., Dumont, D., Berger, S. A., and Ben-David, Y. (2000). Epo regulates erythroid proliferation and differentiation through distinct signaling pathways: implication for erythropoiesis and Friend virus-induced erythroleukemia. Oncogene 19, 2296–2304.

